# Object novelty and value coding segregate across functionally connected brain networks

**DOI:** 10.1101/2019.12.25.887976

**Authors:** Ali Ghazizadeh, MohammadAmin Fakharian, Arash Amini, Whitney Griggs, David A. Leopold, Okihide Hikosaka

## Abstract

Novel and valuable objects are motivationally attractive for animals including primates. However, little is known about how novelty and value processing is organized across the brain. We used fMRI in macaques to map brain activity to fractal patterns varying in either novelty or value dimensions in the context of functionally connected brain networks determined at rest. Results show unique combinations of novelty and value coding across the brain networks. Networks in the ventral temporal cortex and in the parietal cortex showed preferential coding of novelty and value dimensions, respectively, while a wider network composed of temporal and prefrontal areas (TP network), along with functionally connected portions of the striatum, amygdala, and claustrum, responded to both dimensions with similar activation dynamics. Our results support emergence of a common currency signal in the TP network that may underlie the common attitudes toward novel and valuable objects.

## Introduction

Both humans and animals are motivated to interact with novel objects, even if their intrinsic value or utility is unknown ^1–3^. Indeed, novelty preference seems to be genetically hard wired as seen in human infants ^4^ and in lower animals ^5, 6^. One explanation for this behavior could be that novelty carries an inherent value, a value that can be gradually modified in the process of learning ^7^. This prospect is supported by the proposal that reward processing areas are engaged in the processing of novel stimuli ^8, 9 10^. These findings suggest that novelty and value maybe used as a common currency by particular brain networks for goal directed behavior ^11^.

Despite this behavioral and neural link, research that have looked at processing of object novelty and value have traditionally focused on somewhat separate regions (e.g. perirhinal, temporal and prefrontal cortex for novelty ^12–15^ and orbitofrontal cortex and ventral striatum for value ^16–18)^. Hence, a holistic view of the degree of overlap or specialization across the brain for encoding object novelty and value is currently missing. Subjective experience suggests that the distinction between novel objects and valuable objects is not lost to the individual, yet both dimensions seem to orchestrate similar sets of behaviors. A comparative and whole brain mapping of both dimensions can reveal the neural mechanism for common and dissociable processing of object novelty and value.

Notably, for a complex system such as the brain with a patchwork of interconnected regions, network level analyses have become increasingly useful for understanding how functionally connected areas engender, and constrain, neural processing and cognitive functions ^19^. In particular graph theoretical approaches based on resting state fMRI (rfMRI) have been quite successful in revealing behaviorally relevant functional networks across the brain ^20^ with implications for health and disease. Thus, it would be reasonable to ask whether functional networks have relevance for determining an area’s role in encoding novelty and value dimensions.

The present study used fMRI in macaque monkeys to measure brain-wide activations during novel vs familiar (NF) object presentations as well as high vs low value (good/bad or GB) object presentations. The experimental design created novelty and value as orthogonal dimensions: NF objects varied in their perceptual exposure but had no physical reward history, whereas GB objects had equal perceptual exposure but varied in their reward history prior to scanning. During the fMRI scanning sessions, the GB and NF objects were viewed passively. This design ensured that novelty dimension could be compared with value dimension uninterrupted by further reward learning or reward expectation during the scan.

The results revealed the organization of object novelty and value processing across the brain as a gradient from differential to joint processing of each dimension. rfMRI connectivity was utilized to identify and parse functionally coherent networks using graph theoretical approaches. Notably, a core of interconnected temporal and prefrontal cortical areas recently shown to encode long-term value memories ^21^ was found to respond equally to novelty, whereas other networks in ventral temporal and parietal cortex responded preferentially to novelty and value, respectively.

## Results

In order to study the neural correlates of object novelty and value, we performed whole-brain fMRI imaging with two rhesus monkeys (U and D), while exposing each monkey in separate scans to either familiar vs. novel objects or good vs. bad objects. Scans were performed using the contrast agent monocrystalline iron oxide nanoparticles (MION), which, through its isolation and enhancement of local cerebral blood volume, leads to an improved signal-to-noise ratio compared with BOLD (blood-oxygenation-level-dependent)^22^. The objects were computer-generated fractal images, for which familiarity or value was established prior to the fMRI scans. The fMRI scans themselves always involved passive viewing with reward delivered only for maintaining central fixation at random intervals in all blocks. The value dimension was created prior to the scans by associating reward magnitudes with a large number of fractals (>100). These were divided into two groups, corresponding to large and small juice rewards. The monkeys learned these values, thereby creating “good” and “bad” object categories (G and B objects), in the context of a task that involved making saccades to and fixating the fractals shown in the periphery (Fig 1a). To create stable values resistant to forgetting or extinction, such training was repeated for at least 10 days for each fractal prior to the scans ^3, 23^. For the familiarity-novelty dimension, monkeys passively viewed a different group of 8 fractals for more that 10 days, creating a set of long-term familiar objects (F objects, see Methods). There was no reward associated with fractals during the process of perceptual familiarization (Fig. 1b). Novel fractal images (N objects) were created later and were seen only during a single fMRI run. Each monkey saw a large number of novel objects (>160) along with the familiar set during the scans (examples of objects in each dimension shown in Fig 1c). The large number of fractals used and their random assignment to good/bad and novel/familiar groups ensured that any observed fMRI response differences could not be attributed to idiosyncratic features in the stimuli themselves.

**Figure 1:**
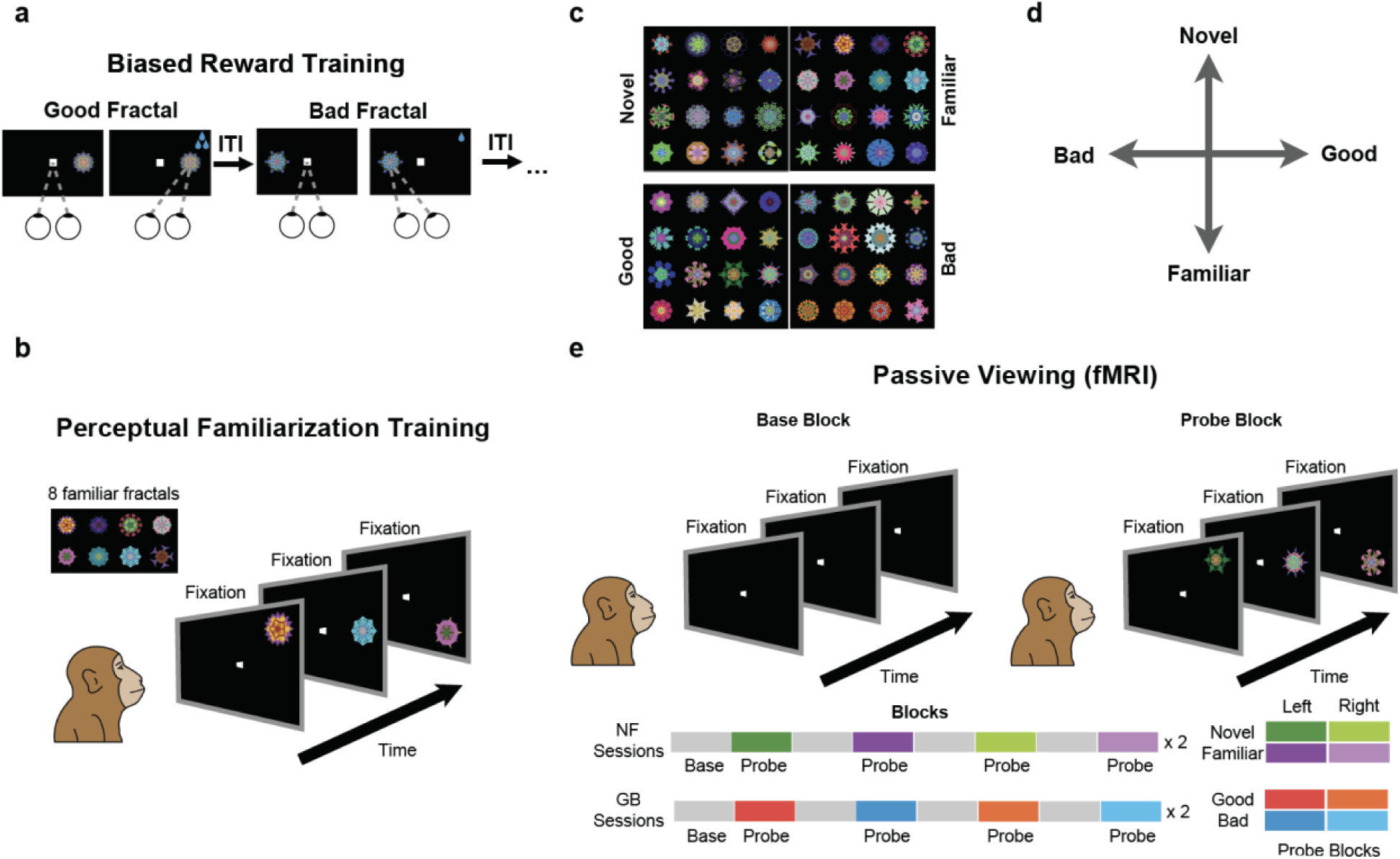
Stimuli and experimental paradigms. **a,** Value training sessions included repeated association of abstract fractal objects with low or high rewards (good and bad fractals respectively) for >10 days. **b,** Perceptual familiarization sessions included repeated exposure of abstract fractal objects in passive viewing and in the absence of reward. **c,** Example fractals used as good/bad and novel/familiar categories. **d,** Schematic of independent novelty and value dimensions. **e,** Test of GB and NF coding in fMRI in a passive viewing task using a block design. In all blocks, subject kept central fixation. In the base blocks, no object was shown. In the probe blocks, objects from one category (good/bad in GB scans and novel/familiar in NF scans, pseudo-randomly ordered through sessions) were shown on the left or right hemifield at 6° eccentricity.

Notably, the good and bad objects had differential reward association but no novelty (equal exposure during the training and test) and the familiar and novel objects had differential perceptual exposure but no history of reward. Thus, past training experience created two distinct dimensions with behavioral significance for the animal objects (perceptual axis and value axis, Fig. 1d). The behavioral significance of each dimension was confirmed by the strong free viewing bias toward good and novel objects The pattern of the free viewing bias supported the cognitive distinctiveness of the two dimensions: the good bias was sustained across repeated exposures while the novelty bias was transient and diminished as novel objects became more familiar with repeated free viewing trials (supp Fig 1).

We mapped the differential responses to novel vs familiar objects (NF scans) and to good vs bad objects (GB scans), performed within a few days of value training or familiarization procedure. The monkey’s task during the fMRI experiment was to simply fixate a white center dot (Fig. 1e). A scanning run consisted of alternating base and probe blocks, each lasting for 30 sec (total of 16 blocks). During the base blocks, only a white fixation dot was shown. During the probe block, the previously experienced fractal objects were presented one at a time in the periphery along with the central fixation. The probe blocks consisted of either good or bad objects in separate blocks in GB scans and novel or familiar objects in separate blocks in NF scans. Objects were presented in the left or the right visual hemifields resulting in four different probe blocks in a given scan (Fig. 1d bottom). These four probe block types were presented in a pseudorandom order.

All fMRI tests were carried out using passive viewing with reward delivered at random time intervals for the successful maintenance of central fixation. The total number of rewards was similar across object types and tasks in both monkeys (supp Fig 2 for NF scans and supp Fig 1 in ^21^ for GB scans). Both animals had extensive experience with the passive viewing task, both outside and inside the scanner (>3 months), before the actual scans. As a result, fixation breaks were infrequent during the scans (<12% fix-break/object) and were not significantly different for object types for both monkeys. Thus, differences in activations between object types were attributed to modulation caused by previous reward or perceptual history. The number of rewards and fixation breaks were also comparable for the left- and right-presentation blocks (supp Fig 2 for NF scans and supp Fig 1 in ^21^ for GB scans). Thus, the interaction between object type and spatial coding were not attributable to differences in reward or fixation breaks between the two visual hemifields during the fMRI tests.

**Figure 2:**
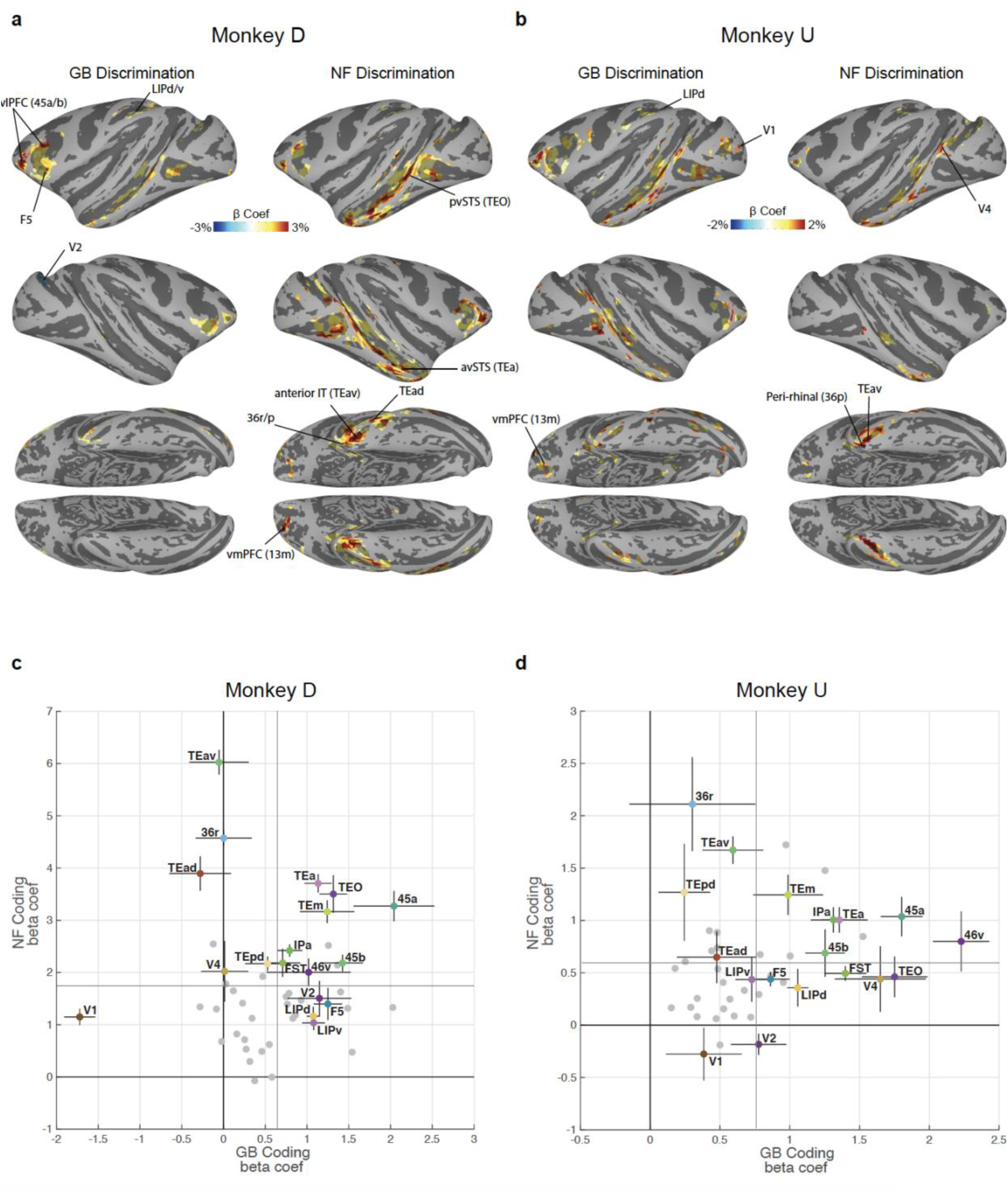
Cortical regions with significant novelty and value coding. **a,** Cortical regions’ beta coefficients in significantly responsive voxels for GB contrast (left: value coding) and NF contrast (right: novelty coding) in monkey D shown in standard space ^41^ (*P <* 0.001*, α <* 0.01 cluster corrected). **b,** Same format as **a** but for Monkey U (*P <* 0.001*, α <* 0.01 cluster corrected). **c,** Average GB (horizontal axis) and NF (vertical axis) beta coefficients for top 10 visually activated voxels across the entirety of a cortical area with at least one cluster of activated voxels in either GB or NF scans in monkey D. Each point represents one cortical region. A number of key areas are annotated. The vertical and horizontal thin gray lines specify mean values of NF coefficients and GB coefficients across all activated cortical regions, respectively. The errorbars indicate the s.e.m. across novelty and value dimensions. **d,** Same format as **c** but for monkey U.

### Cortical regions sensitive to novelty and value

Cortical areas showing robust differentiation of NF and GB are illustrated in Fig. 2. As previously described ^21^, results from GB scans showed robust discrimination in several cortical regions most prominently in posterior ventral superior temporal sulcus (pvSTS) including areas such as TEO, FST and TE and in ventrolateral prefrontal cortex (vlPFC) including areas 45 and 46 (Figs 2a-b) in both monkeys. Interestingly, NF scans showed robust novelty-familiarity discrimination in many of the same areas activated by value discrimination (including pvSTS and vlPFC). Despite the overall similarity, NF and GB coding across the brain showed some notable differences. For example, the activation in STS seemed to include more anterior regions (avSTS) in NF vs GB scans (Figs 2a-b). This difference was particularly evident in stronger and broader activity in the anterior IT and perirhinal cortex (TEav/d, 35, 36r/p/c) in NF vs GB scans. A quantitative comparison of activity in the posterior vSTS (TEO, FST, IPa, V4, MT) with anterior vSTS (TEm/a) confirms stronger NF vs GB coding in anterior vs posterior vSTS (supp Fig 3). Finally, area LIP in parietal cortex was more activated in GB vs NF scans in both monkeys (Fig 2a-b).

**Figure 3:**
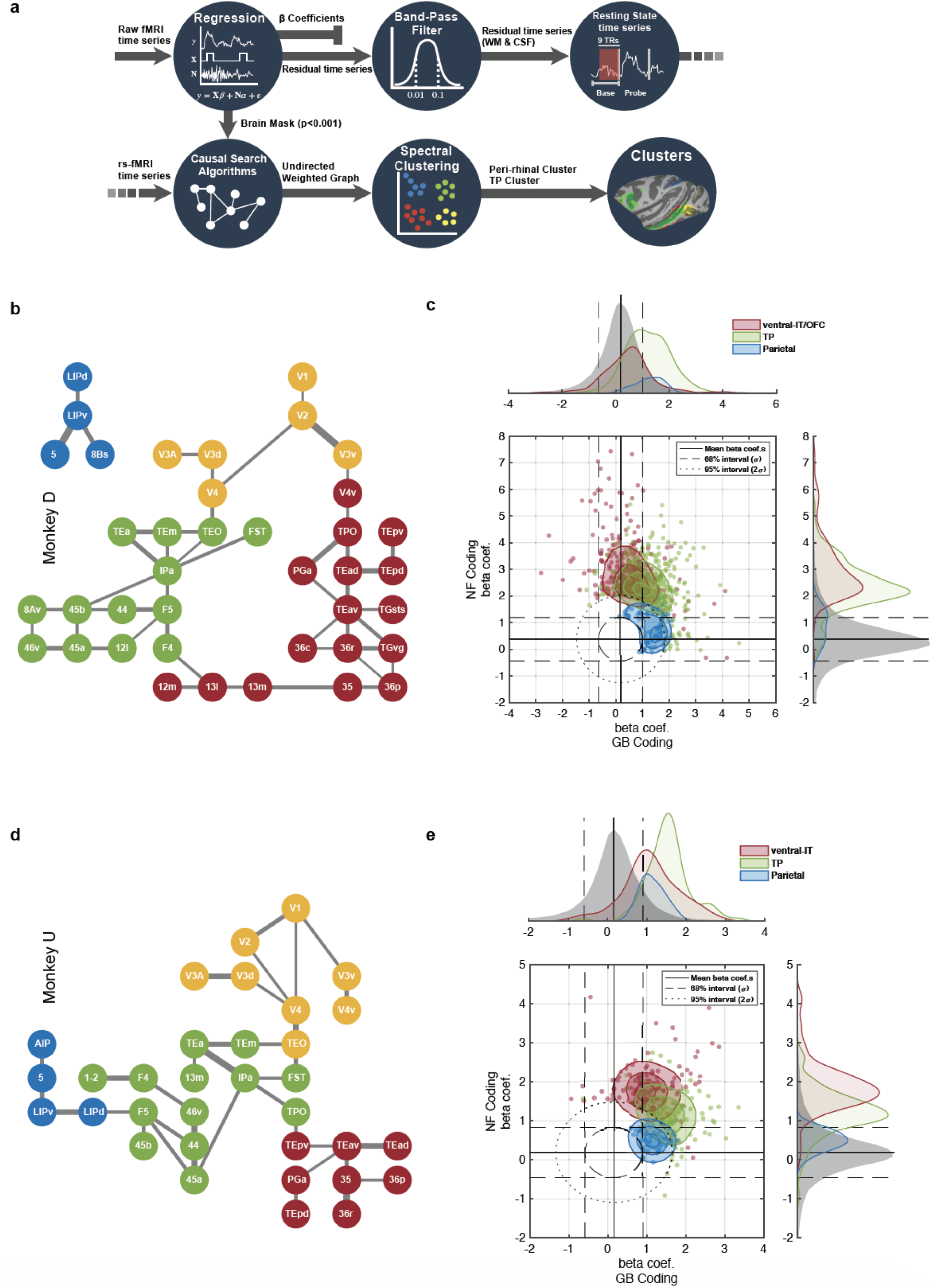
Resting state cortical networks account for novelty and value coding for member areas. **a,** Schematic of the functional connectivity analysis steps based on resting correlation. The first base blocks from all of the runs were selected and concatenated after regression of nuisance parameters and band-pass filtering. The representative graph is extracted via PC-stable algorithm (see methods). Having estimated the weighted undirected graph, spectral clustering was used to partition the graph into four networks. **b,** Functional connectivity graph segmented into 4 networks using unsupervised spectral clustering in monkey D. Occipital: yellow, TP: green, ventral-IT: red and parietal: blue **c,** The NF/GB distribution of beta coefficients in all subcortical and cortical areas (whether or not significant) was considered jointly as the null distribution and iso-probability contours marking the 68% (dashed line) and 95% (dotted line) interval of the joint distribution were drawn. Cortical voxels falling outside these confidence intervals were plotted and colored according to network membership (95% used for ventral-IT and TP networks and 68% used for parietal network). The marginal distributions of beta coefficients for the null distribution and for the three networks are also depicted across the GB and NF axes. (**d**-**e**) same format for monkey U.

There were also some differences between the monkeys. In monkey D the OFC (areas 13m/l) was more active in NF discrimination task, while in monkey U, GB discrimination was more prominent in this region. Also, the location of activity in the occipital areas was inconsistent between the two monkeys (Fig 2). Finally, in monkey D the right V1-V2 cortexes had negative GB coefficient with right lunate sulcus reaching significance. This was the only negative GB coding found across the cortex in either monkey (Fig. 2). The complete list of significantly activated cortical regions in NF and GB scans in each monkey is reported in Table 1 and anatomical annotations are shown in supp Figure 4.

**Figure 4:**
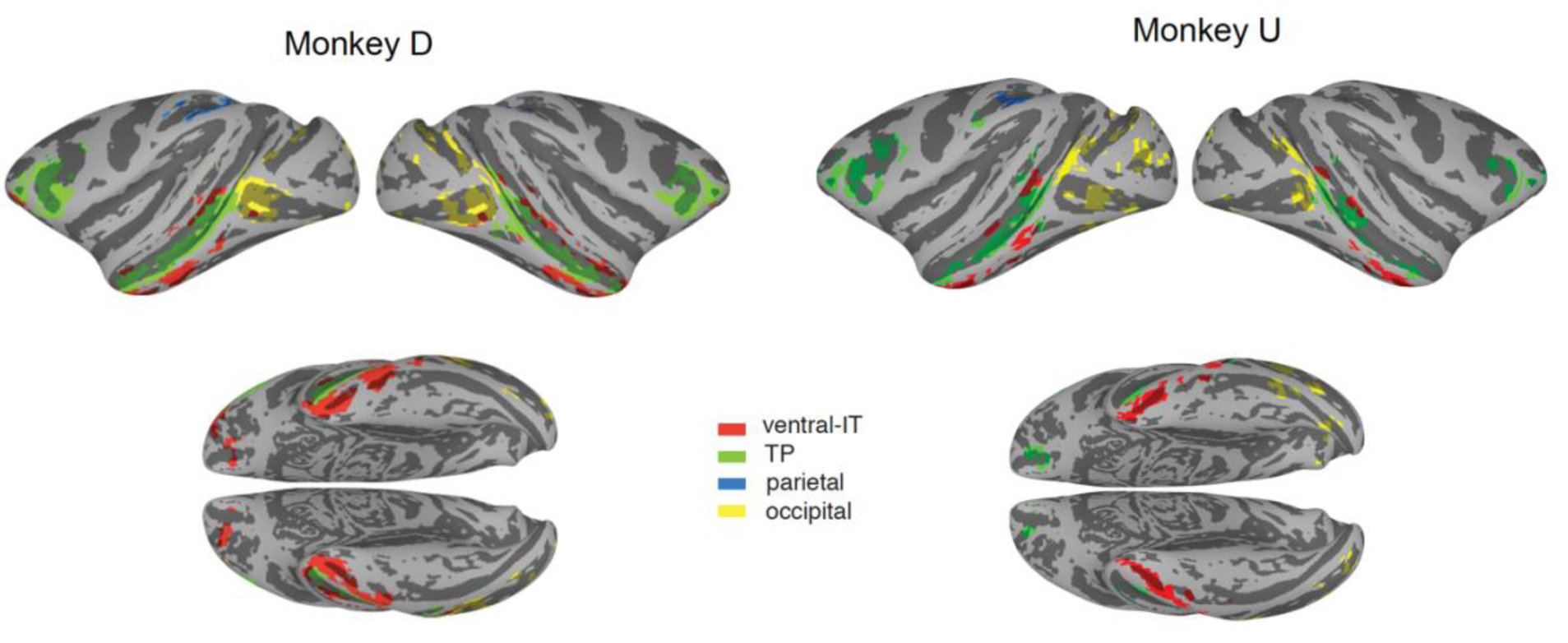
Localization of the four resting state cortical networks. The four networks for monkey D (left) and monkey U (right) color-coded and shown on standard brain surface ^41^.

**Table 1:**
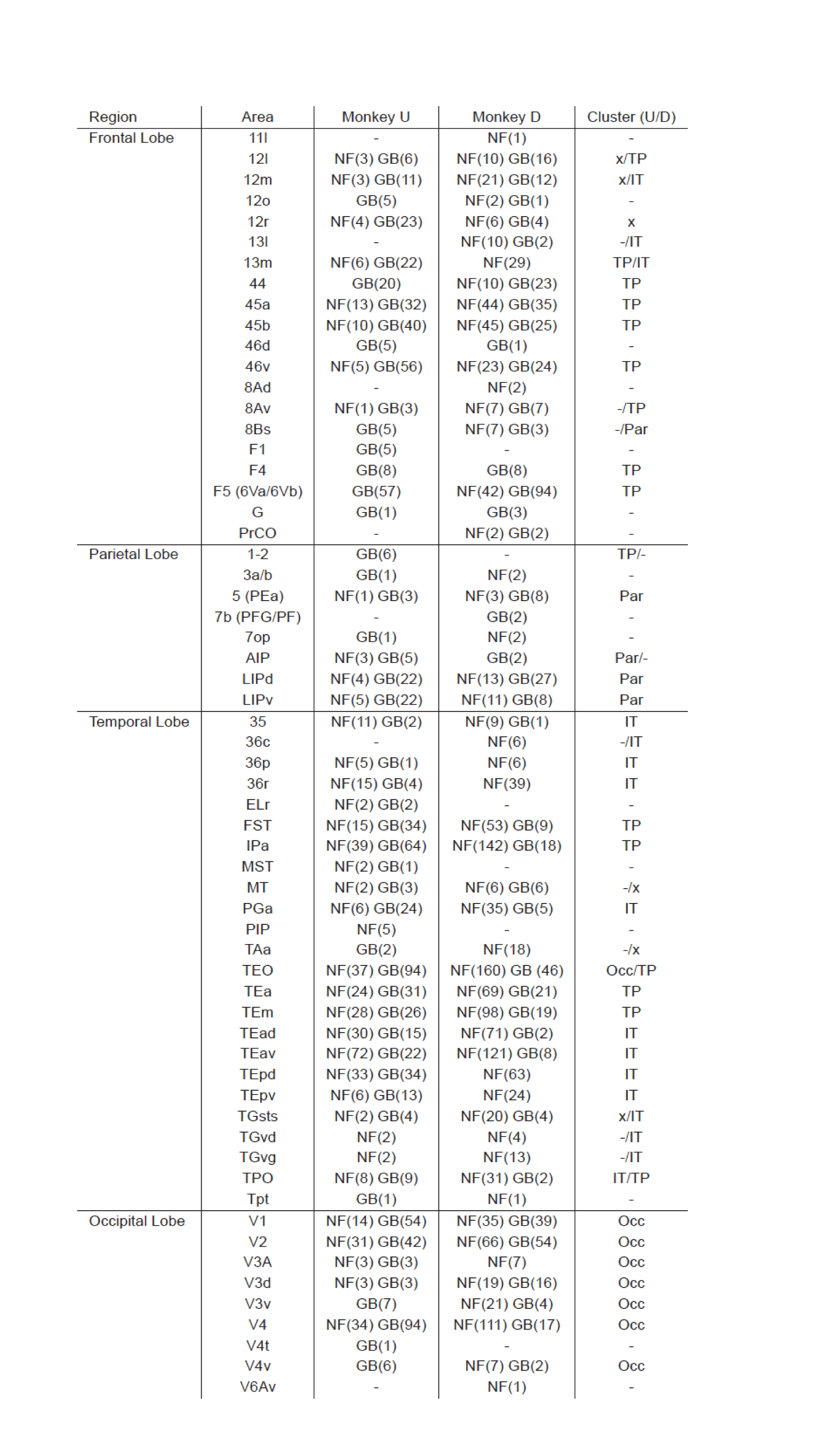
List of cortical ROIs active in NF and GB task for both monkeys. Values in the parentheses are number of voxels with *P <* 0.001 which are cluster-corrected for the whole brain at *α <* 0.01. The rightmost column specifies the cluster each region belongs to for monkeys U and D. (-: omitted, x: not in any cluster).

| Region | Area | Monkey U | Monkey D | Cluster (U/D) |
| --- | --- | --- | --- | --- |
| Frontal Lobe | 11l | - | NF(1) | - |
|  | 12l | NF(3) GB(6) | NF(10) GB(16) | x/TP |
|  | 12m | NF(3) GB(11) | NF(21) GB(12) | x/IT |
|  | 12o | GB(5) | NF(2) GB(1) | - |
|  | 12r | NF(4) GB(23) | NF(6) GB(4) | x |
|  | 13l | - | NF(10) GB(2) | -/IT |
|  | 13m | NF(6) GB(22) | NF(29) | TP/IT |
|  | 44 | GB(20) | NF(10) GB(23) | TP |
|  | 45a | NF(13) GB(32) | NF(44) GB(35) | TP |
|  | 45b | NF(10) GB(40) | NF(45) GB(25) | TP |
|  | 46d | GB(5) | GB(1) | - |
|  | 46v | NF(5) GB(56) | NF(23) GB(24) | TP |
|  | 8Ad | - | NF(2) | - |
|  | 8Av | NF(1) GB(3) | NF(7) GB(7) | -/TP |
|  | 8Bs | GB(5) | NF(7) GB(3) | -/Par |
|  | F1 | GB(5) | - | - |
|  | F4 | GB(8) | GB(8) | TP |
|  | F5 (6Va/6Vb) | GB(57) | NF(42) GB(94) | TP |
|  | G | GB(1) | GB(3) | - |
|  | PrCO | - | NF(2) GB(2) | - |
| Parietal Lobe | 1-2 | GB(6) | - | TP/- |
|  | 3a/b | GB(1) | NF(2) | - |
|  | 5 (PEa) | NF(1) GB(3) | NF(3) GB(8) | Par |
|  | 7b (PFG/PF) | - | GB(2) | - |
|  | 7op | GB(1) | NF(2) | - |
|  | AIP | NF(3) GB(5) | GB(2) | Par/- |
|  | LIPd | NF(4) GB(22) | NF(13) GB(27) | Par |
|  | LIPv | NF(5) GB(22) | NF(11) GB(8) | Par |
| Temporal Lobe | 35 | NF(11) GB(2) | NF(9) GB(1) | IT |
|  | 36c | - | NF(6) | -/IT |
|  | 36p | NF(5) GB(1) | NF(6) | IT |
|  | 36r | NF(15) GB(4) | NF(39) | IT |
|  | ELr | NF(2) GB(2) | - | - |
|  | FST | NF(15) GB(34) | NF(53) GB(9) | TP |
|  | IPa | NF(39) GB(64) | NF(142) GB(18) | TP |
|  | MST | NF(2) GB(1) | - | - |
|  | MT | NF(2) GB(3) | NF(6) GB(6) | -/x |
|  | PGa | NF(6) GB(24) | NF(35) GB(5) | IT |
|  | PIP | NF(5) | - | - |
|  | TAa | GB(2) | NF(18) | -/x |
|  | TEO | NF(37) GB(94) | NF(160) GB (46) | Occ/TP |
|  | TEa | NF(24) GB(31) | NF(69) GB(21) | TP |
|  | TEm | NF(28) GB(26) | NF(98) GB(19) | TP |
|  | TEad | NF(30) GB(15) | NF(71) GB(2) | IT |
|  | TEav | NF(72) GB(22) | NF(121) GB(8) | IT |
|  | TEpd | NF(33) GB(34) | NF(63) | IT |
|  | TEpv | NF(6) GB(13) | NF(24) | IT |
|  | TGsts | NF(2) GB(4) | NF(20) GB(4) | x/IT |
|  | TGvd | NF(2) | NF(4) | -/IT |
|  | TGvg | NF(2) | NF(13) | -/IT |
|  | TPO | NF(8) GB(9) | NF(31) GB(2) | IT/TP |
|  | Tpt | GB(1) | NF(1) | - |
| Occipital Lobe | V1 | NF(14) GB(54) | NF(35) GB(39) | Occ |
|  | V2 | NF(31) GB(42) | NF(66) GB(54) | Occ |
|  | V3A | NF(3) GB(3) | NF(7) | Occ |
|  | V3d | NF(3) GB(3) | NF(19) GB(16) | Occ |
|  | V3v | GB(7) | NF(21) GB(4) | Occ |
|  | V4 | NF(34) GB(94) | NF(111) GB(17) | Occ |
|  | V4t | GB(1) | - | - |
|  | V4v | GB(6) | NF(7) GB(2) | Occ |
|  | V6Av | - | NF(1) | - |

We then leveraged the independence of two task contexts (Fig. 1d) to represent the average NF and GB beta coefficients in a two-dimensional plot for all anatomical areas activated in either scan type (Fig. 2c-d). To avoid selection bias, beta coefficients were averaged for the 10 most visually active voxels (beta coefficients in probe vs base contrast) in each anatomical area, regardless of NF or GB coding strengths. The vlPFC (e.g. 46v, 45a/b) and vSTS regions (e.g. IPa, TEa/m) showed larger than average NF and GB coding in both monkeys among regions with significant coding in either dimension. On the other hand, ventral IT including areas TEav/d, TEpd and perirhinal cortex (36r/p) had larger than average NF beta coefficients and below average GB beta coefficients in both monkeys. The GB discrimination was small in monkey U and almost absent in monkey D in ventral IT regions. Finally, parietal areas including area LIPd showed below average NF discrimination but better than average GB discrimination.

These findings show a gradient among regions in their sensitivity to object novelty and value with areas close to the cardinal axes showing preferential coding of either novelty or value dimensions while others falling in the upper-right quadrant exhibiting joint representation of novelty and value in support of the common currency theorem.

### Resting state networks among novelty and value-coding cortical areas

The gradient of GB and NF coding across cortical regions suggest the existence of separate functional networks for processing of object attributes. We examined this possibility by quantifying the functional connectivity (FC) between activated regions using the resting block data in the beginning of each scan. Briefly, preprocessing steps for resting state correlation including nuisance regression, bandpass filtering and white-matter and ventricle signal cancellation was done on the first resting block in each scan as shown in Figure 3a. Pairwise correlations based on spontaneous activity of all activated regions were used to arrive at a graphical representation of connectivity using a causal search algorithm (PC-stable ^24^, see methods for details). Briefly, the PC-stable algorithm is a constraint-based algorithm that aims to extract the causal graph skeleton based on conditional independence tests (using partial correlations). Starting with the fully connected graph, the algorithm removes the edges between nodes that end up with non-significant partial correlation given all possible combinations of neighboring node subsets. For the remaining edges, in our implementation, the weights were set to the lowest significant partial correlation between node pairs representing the net correlation between two nodes. The resulting graph is illustrated in Fig. 3b, d for monkey D and U, respectively in which anatomical areas with significant NF or GB coding are represented as graph nodes. Notably and despite the fact that the causal search algorithm had no prior knowledge of anatomical connections between the nodes, the resulting graph conformed well with the known hierarchy of visual information starting from V1 and moving successively across areas in the ventral stream from the posterior to anterior IT cortex to finally arrive in prefrontal cortex ^25^.

The pattern and strength of connectivity between nodes (cortical areas) was used to divide the cortex into four clusters or networks. Clustering of graph nodes, was done by first representing the nodes in the eigenvector space of graph Laplacian matrix and then using a clustering algorithm (k-means) to group the nodes (spectral clustering). Interestingly, this approach delineated similar networks in both monkeys namely: 1) an occipital network including early visual areas V1-V4, 2) a vSTS and vLPFC network including areas FST, IPa, TEa/m and areas 45a/b, 46 which was previously found using a different analysis technique and was referred to as the TP network ^21^ 3) a ventral-IT network including areas TEav/d and perirhinal cortex (areas 36r/p and 35) and 4) a parietal network including areas LIPd and LIPv (Fig 3b,d).

Besides the overall similarity, there were some variations between the networks in two monkeys. First in monkey D, two branches stemmed from the visual cortex; one of which led to the TP network and the other to the ventral-IT network, while in monkey U the connectivity flow to ventral-IT network was via the TP network. Second, OFC area 13m was grouped with ventral-IT (network 3) in monkey D and with TP network in monkey U. Third, the parietal network was stand alone in monkey D but was connected to the prefrontal part of TP network in monkey U. Despite the observed differences, the overlay of the 4 cortical networks showed a qualitatively similar segmentation of the cerebral cortex in both monkeys (Fig 4).

### Segregation of novelty and value coding by resting state networks

Next, we wanted to know if the detected networks could account for the differences in novelty and value coding across the cortical areas and between subjects. To do so, we plotted the NF vs GB beta coefficients for all activated voxels in TP network, ventral-IT network and parietal network (Fig 3c,e). In addition, the joint distribution of NF and GB beta coefficients for all voxels across all the gray matter (cortical and subcortical areas) was constructed as the 2D null distribution in this space. This allowed us to use equiprobability confidence ellipses using the null distribution to determine the voxels that show strong novelty or value coding outside the confidence intervals. Figure 3c,e shows voxels laying outside the confidence ellipses, colored according to their resting state network membership. Majority of voxels in parietal network did not have strong GB or NF coding that fell outside the 95% confidence ellipse. To visualize the relative coding of this network, we lowered the confidence threshold for this network to 68% while for the other two networks 95% confidence ellipse was used. Despite, some differences between the two monkeys, the relative spatial arrangement of voxels in the NF-GB coordinate system for the three networks were strikingly similar. In both monkeys, NF coding showed a gradient from high to low going from ventral-IT to TP to parietal networks. TP network showed strong co-coding of novelty and value while ventral-IT and parietal cortex were more biased toward novelty and value coding, respectively (Fig 3c,e). Unlike the other 3 resting state networks, the occipital network results were not well localized in the NF-GB coordinate space (supp Fig 5) which may reflect the inconsistent and variable beta coefficients in the occipital regions across the two monkeys (Fig 2). The similarity of atlas-based results in Figure 2 and data-driven FC results in Figure 3c,e is striking since there is no particular reason the FC networks should abide by abide by cytoarchitectonic boundaries (Fig 4 networks vs supp Fig 4).

**Figure 5:**
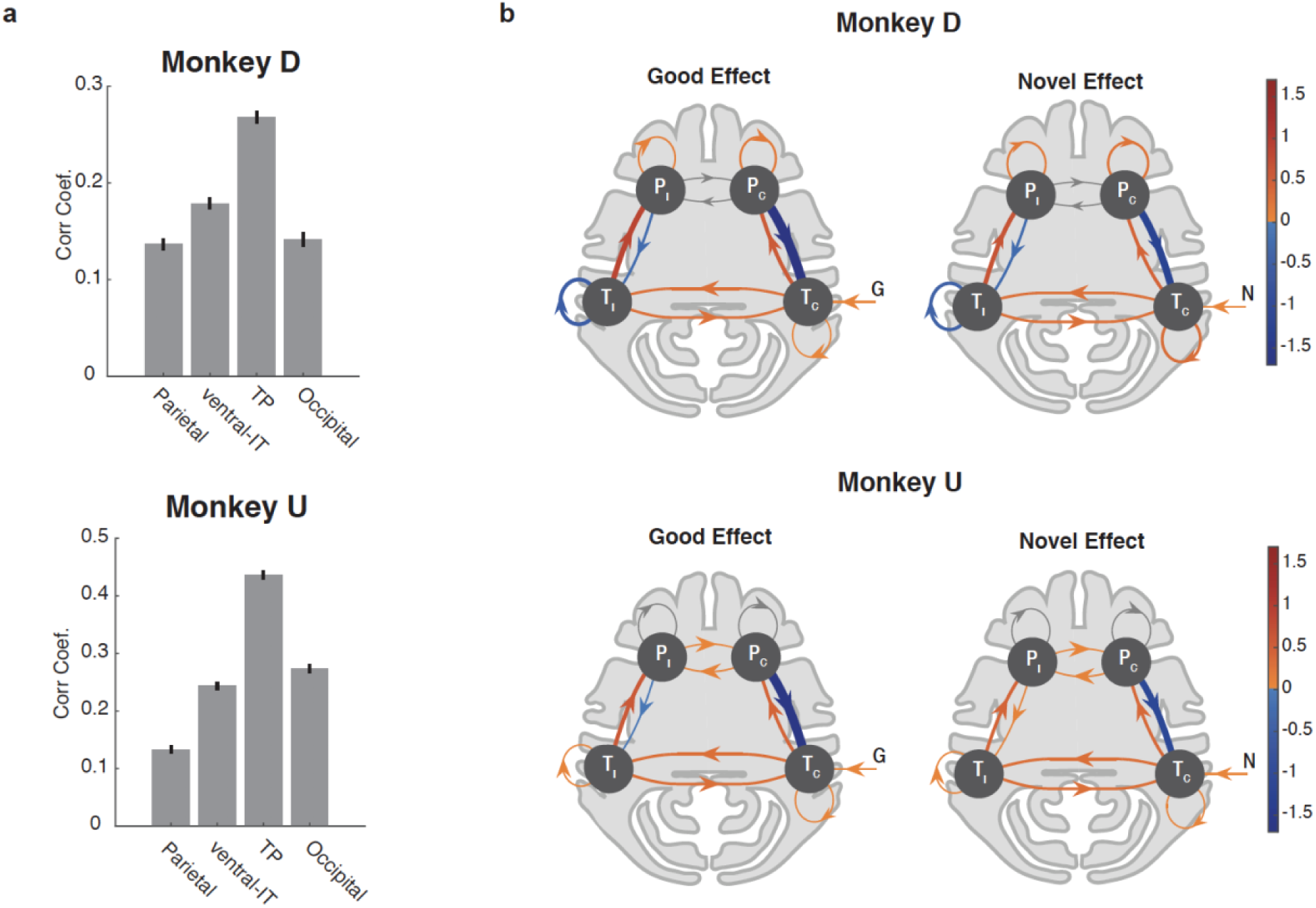
Similarity of time-course and dynamics of NF/GB activations in the TP network. **a,** Average MION signal correlation coefficient between NF and GB time-courses in voxels in each network. Error bars indicate s.e.m across voxels. **b,** DCM model of the prefrontal-temporal interaction during good and novel object presentation. Bayesian model selection was used to arrive at the best DCM model for each monkey. For both monkeys the best model had contralateral visual input to the temporal node. The self-connections were modulated by input type (G,B,N,F) in monkey D but not in monkey U (see supp Fig 7 for details). The edge weights for presentation of good (left) and novel (right) were combined (averaged) across right and left blocks and are presented in contra vs ipsi format (T_c,_ T_i_ : contra and ipsilateral temporal nodes, P_c,_ P_i_ : contra and ipsilateral prefrontal nodes). The line widths are scaled by the absolute value of the edges. Signed weights are color coded. For Monkey U self-connections in P nodes were inconsistent between the two hemispheres and are grayed out. Monkey D: T node included TEa, TEm, TEO, FST, IPa and P node included 8Av, 45b, 44, F5, F4, 12l, 45a, 46v. Monkey U: T node included TEa, TEm, TPO, FST, IPa and P node included 13m, 45b, 44, F5, F4, 45a, 46v.

These findings suggest that network membership for an area can have consequences for its NF or GB coding. Thus, differences in network membership for a given area across subjects could be predictive of different roles in NF and GB coding for that area. Interestingly, this seems to be the case for the OFC. In monkey D, the OFC (13m/l) had better NF coding but weaker GB coding and in this monkey, OFC was a part of ventral IT network (Fig 3b). In monkey U, the OFC (13m/l) had good GB and NF coding and here, it belonged to the TP network based on resting connectivity (Fig 3d). Furthermore, for monkey U whose ventral-IT network had larger GB-coding in comparison to monkey D, ventral IT network had connectivity to visual cortexes via the TP network which encoded both NF and GB dimensions.

### Similarity of novelty and value coding dynamics in the TP network

The fact that TP network showed strong activation to both object novelty and value, suggests that this network may treat object novelty and value similarly and for similar purposes. To further test this idea, we looked at the temporal dynamics of GB and NF signaling in the TP network. To this end, we first did a simple correlation between the time course of activity during the NF scans and GB scans in the TP networks as well as in the other functional networks. The idea was that if TP network treats novelty and value dimensions similarly, the time series of activations to NF and GB scans, should have a higher correlation in this network compared to other networks. The results showed higher correlation in the TP network compared to other networks (Fig 5a). Notably, this effect was not due to higher visual activity in TP compared to other networks since TP visual activation was not necessarily higher than other networks (supp Fig. 6a) and furthermore a positive relation between NF/GB timeseries correlation and visual activation across voxels was not found (supp Fig. 6b).

**Figure 6:**
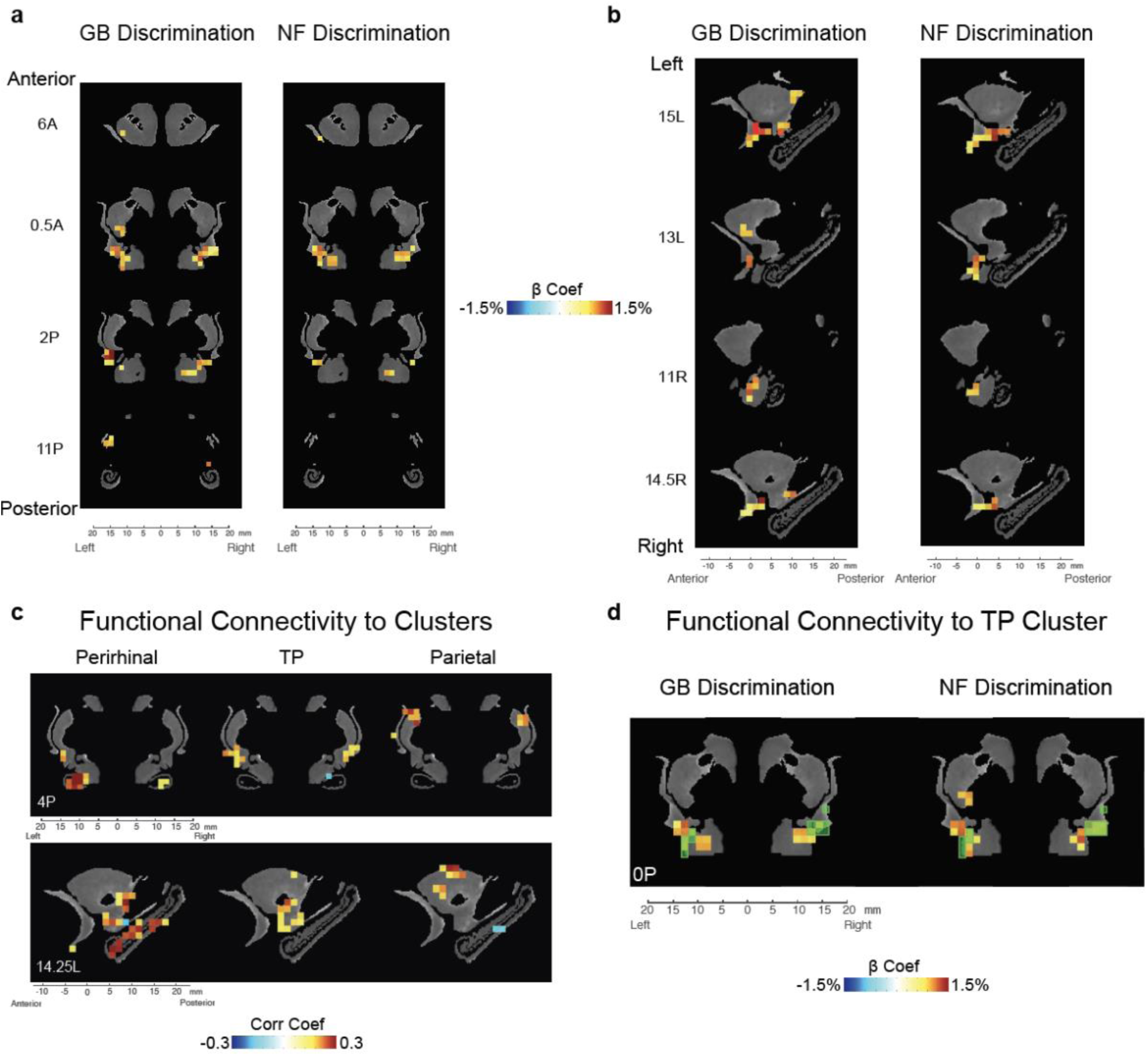
Subcortical novelty and value coding in striatum, amygdala, claustrum, and hippocampus. **a,** GB (left) and NF (right) significantly active voxels in coronal view (P<0.001, α<0.01 cluster-corrected). **b,** Same as **a** in sagittal view. **c,** Voxels with significant resting correlation with ventral-IT, TP, and parietal networks in coronal (top) and sagittal (bottom) views (P<0.001, α<0.01 cluster-corrected). **d,** Overlay of TP-connected voxels (green transparent squares) and voxels with significant GB (left) and NF (right) coding (beta coefficients of GB and NF shown). Data in this figure are from monkey U.

To compare the temporal activation dynamics between NF and GB in the TP network more precisely, dynamic causal modeling (DCM ^26^) was used. The proposed DCM model included one node in the temporal and prefrontal cortex in each hemisphere (a total of 4 nodes, Fig 5). Model fitting was done for multiple plausible architectures (supp Fig 7a-b). Figure 5b shows the architecture with the highest posterior probability. In this architecture, the temporal node in each hemisphere receives direct visual information from the contralateral visual hemifield. Interestingly, the dynamics of activations for good and novel objects turned out to be similar within each monkey as evident by the pattern of weights between the nodes (Fig 5b and supp Fig 7c). The same was true for the dynamics of activations for bad and familiar objects (supp Fig 7d-e). This further supports the hypothesis that TP networks treats perceptual and value dimension as interchangeable dimensions.

The DCM model also revealed the likely mechanism for the interaction of prefrontal and temporal nodes in the TP network for coding of GB and NF dimensions. In both monkeys, a negative feedback from vlPFC to vSTS was found in both NF and GB scans. Such inhibitory top-down influence was observed despite the fact that negative self-feedback in vSTS and vLPFC was allowed in the model (thus ruling out simple model stabilization mechanisms). Such negative feedback represents a top down mechanism from higher level cortical areas (vlPFC) to control the level of activation to good and novel signals in lower level sensory areas in (vSTS). Together, these results suggest that up to the fMRI temporal resolution the perceptual and value dimensions are being coded similarly in the TP network.

### Subcortical substrates for novelty and value coding

Previously, it was shown that value memory is well represented in certain subcortical areas including ventral claustrum, ventral putamen, caudate tail and lateral amygdala ^21^. Notably, these regions were previously found to have strong functional connectivity to the TP network. Given the role of TP in co-coding of value and perceptual memory, we hypothesized that the same subcortical regions should also show GB and NF discriminations. This was indeed the case. As can be seen in Figure 6a-b for monkey U, the aforementioned subcortical regions in claustrum, striatum and amygdala also showed significant NF coding. In the striatum, the GB and NF coding areas were localized in the caudal and ventral part which are called the tail of the caudate nucleus (CDt) ^27^ and the tail of the putamen (PUTt) ^28^. Figure 6c shows the parcellation of the subcortical regions by their resting state functional connectivity to the TP, ventral-IT and parietal cortical networks. One can see that ventral subcortical areas including the hippocampus and ventral amygdala were connected to the ventral-IT network. The areas connected to the TP network include somewhat more dorsal areas specially in ventral putamen and ventral claustrum while areas connected the parietal network are mostly in the dorsal striatum. This result is largely consistent with known anatomical connections ^14, 29, 30^. The voxels with functional connectivity to TP network showed GB and NF coding (Fig 6d). However, the areas with connectivity to ventral IT such as hippocampus did not show strong NF coding and the areas connected with parietal network did not show strong GB coding. Thus, despite having significant resting correlation, at least in our task, the ventral-IT and parietal networks did not seem to share the value and perceptual memory information with their functionally connected subcortical areas. Similar results for subcortical areas was observed in monkey D (supp Fig 8). In this monkey, GB coding in subcortical areas was somewhat weaker and NF coding was observed in a few voxels in hippocampus. The complete list of significantly activated subcortical regions in NF and GB scans for both monkeys is reported in Table 2.

**Table 2:** List of subcortical ROIs active in NF and GB task for both m onkeys. Values in the parentheses are number of voxels with *P <* 0.001 in each area with small area cluster correction at *α <* 0.01. The voxels in hippocampus of monkey D belonged to the most posterior part as depicted in sup Fig 8b.

| Area | Monkey U | Monkey D |
| --- | --- | --- |
| Striatum | NF(12) GB(41) | NF(31) GB(6) |
| Clastrum | NF(22) GB(29) | NF(43) GB(5) |
| Amygdala | NF(13) GB(17) | NF(33) GB(4) |
| Hippocampus | - | NF(4) |

## Discussion

Novelty seeking guides decisions when interacting with unknown objects. On the other hand, when dealing with familiar objects value seem to be a key driving force for decision making and choice. The aim of this study was to reveal the organization of object novelty and value processing across the brain. Our results show widespread cortical activation in both NF and GB discrimination in all four brain lobes with both overlapping and non-overlapping regions (Fig 2 and supp Fig 4). Resting state analysis showed that the degree of NF-GB co-coding across different cortical regions could be accounted for by the functional connectivity among the regions. Specifically, the FC between cortical regions with significant NF or GB coding revealed four networks: an occipital network which included early visual areas, a temporal-prefrontal network which included areas in vSTS and vlPFC (the TP network), a ventral-IT network which included TEa and perirhinal cortex and a parietal network which included area LIP (Fig 3-4). Among these functional networks, the TP network was found to be sensitive to both GB and NF dimensions. The ventral-IT network was preferentially sensitive to the NF dimension. The parietal network was preferentially sensitive to the GB dimension (Fig 3c,e). For the TP network, the activation dynamics and interplay of the temporal and prefrontal nodes in coding object novelty and value were also similar (Fig 5 and supp Fig 7). Notably, the subcortical areas with strong connectivity to TP but not the other two networks also showed co-coding of NF and GB dimensions (Fig 6).

The TP network was previously found to represent learned object values in long-term memory ^21^. The co-activation of this network to object novelty suggests that novelty maybe processed as an intrinsically valuable property in the brain (common currency theorem). On the other hand, areas in ventral IT including TEav and TEad and perirhinal areas such as area 36r showed much stronger sensitivity to NF dimension. This result is consistent with the role of medial temporal lobe (MTL) in recognition memory ^14^. It is thus possible that the novelty is first detected in the ventral-IT network and then transmitted to the TP network. Also interesting was the fact that the parietal areas including area LIP was more sensitive to object value rather than object novelty. This result is among the few differences reported between LIP and vlPFC for object prioritization. Given the role of parietal and TP networks in attentional modulation, their different coding sensitivity in GB-NF coordinate system indicate that the visual attention mediated by the two networks may be of different types and for different purposes.

Recent studies using monkey electrophysiology have revealed hot spots in basal ganglia including CDt and PUTt to encode object intrinsic values ^28, 31, 32^. Both CDt and PUTt are known to receive direct inputs from vSTS ^29, 33^ and to send signals indirectly to the temporal cortex ^34^ and the prefrontal cortex ^35^ the areas that are found to form the TP network via the caudal-dorsal-lateral part of the substantia nigra reticulata (cdlSNr) ^36^. Given the observed rapid and robust value discrimination in dSNr and the fact that basal ganglia have connections to cortical areas through the thalamus and are shown to lead cortex in value learning ^37^ it is reasonable to assume basal ganglia as the source of value coding for the TP network. Moreover, our fMRI study shows that the PUTt encode both GB and NF dimensions (Fig 6 and supp Fig 8). Previous electrophysiological studies actually showed that CDt neurons responded to novel objects more strongly than to familiar objects ^38^, in addition to their clear value coding ^27, 32^. These data suggest that the GB and NF coding of the TP network may be caused by the inputs from the basal ganglia.

Interestingly, resting state correlations with the three cortical networks revealed different patterns of connectivity to subcortical areas across the dorsoventral axes (Fig 6c and supp Fig 8c). We found co-coding of NF and GB in the ventral claustrum and lateral amygdala both of which showed good connectivity to the TP network during rest (Fig 6c). However, despite the NF coding in ventral-IT network and GB coding in parietal network, the subcortical areas connected to these areas did not show significant NF or GB coding. This result was most striking in the hippocampus that showed strong connectivity to ventral-IT network but almost no significant NF coding (Fig 6, supp Fig 8 and Table 2). While the lack of NF coding in hippocampus may be surprising, it is largely consistent with what is known about the role of hippocampus in recollection rather than recognition of familiar stimuli ^14^.These results suggest that despite the resting connectivity some of the active functionality of cortical networks are not relayed subcortically. The factors that control such information transmission or lack thereof requires future investigations.

There are so many objects in real life, but we (and animals) can choose only a few of them in many cases. We may have experienced many of them previously (familiar objects) and therefore their values may be known already. The others are novel objects whose values are unknown. Then, there are two goal-directed behaviors. The first is to choose a valuable (good) object among familiar objects. The second is to choose and explore novel objects. Such a novelty bias could well be related to the preference for obtaining information about objects ^39^ which even though activates reward circuitry is shown to be independent from reward coding ^9, 40^. Note, that even in a world where many novel objects are aversive, novelty bias may be justified to know what objects to avoid in the future. It is possible that evolution have hard wired the brain to increase exposure to novel objects to learn better models of the environment. The overlapping representation of NF and GB dimensions in the TP network suggests that many aspects of the neural and physiological processing related to object novelty and value that is directed by this network could be similar. Indeed, many of the behavioral outputs toward novel and valuable object are found to be similar ^3, 7^. However, as expected, the source of motivation is not lost in the brain, since there are networks that preferentially encode novelty or value dimensions but not both (Fig 3).

In summary and together with previous evidence, our results suggest that perceptual memory (novelty vs familiarity) that arises from the ventral-IT network and value memory that arises from the basal ganglia are integrated in the TP network. Value and novelty both demand the attention of the animal and promote interaction with the objects. The TP network is involved in both detailed object processing (the T node) and executive planning (the P node). Thus, the common activation in TP for good and novel objects can be key for preferential processing and orientation toward these objects (common currency theorem, supp Fig 1). Future experiments are needed to address the differences in causal role of each regions in specific behaviors evoked by NF and GB dimensions. It would also be interesting to see whether aberrated processing in TP network that affects GB dimension also affect the NF dimension and vice versa.

## SI Methods

### Subjects

Two adult rhesus monkeys (U: female, D: male) were involved in the study. In order to prevent head movements, the monkeys were implanted with a MR compatible head post prior to the training procedure. In addition, a scleral search coil was implanted in one eye of each monkey for eye tracking during training outside the scanner. Inside the scanner, eye position was monitored by an MR compatible infrared camera (MRC camera 60 Hz, SMI IView x 2.6, tracking resolution <0.1°). All animal care and experimental procedures were approved by the National Eye Institute Animal Care and Use Committee and complied with the Public Health Service Policy on the humane care and use of laboratory animals.

### fMRI Scanning

Awake animals sat upright in an MR-compatible chair and viewed visual stimuli via a mirror reflecting a display positioned above their head. A 4.7-T, 60-cm vertical MRI scanner (Bruker Biospec) equipped with a Brucker S380 gradient coil was used to acquire the fMRI data while the monkeys performed the passive viewing task. Functional MRI images were collected using a T2* echo planar images (EPI) sequence with 1.5 mm^3^ isotropic spatial resolution, repetition time (TR) = 2.5s, echo time (TE) = 14ms, and flip angle = 85°. Prior to each scanning session, a T2* contrast agent (MION) was injected to each subject (∼10mg/kg). High-resolution anatomical scans with 0.5mm^3^ isotropic spatial resolution were also collected from anesthetized monkeys by a horizontal 4.7-T MRI (Bruker Biospec 47/60).

### Stable Value Training

The biased reward training consisted of an object-directed saccade task to train the values of each good and bad fractal objects (Fig. 1a). Each training block of this task used a set of 8 fractals (4 Good and 4 Bad fractals). Following the subject fixating on a central white dot, an object appeared from the set at one of the 8 peripheral locations (15° eccentricity). After a 400 ms period, the fixation dot disappeared and the subjects were supposed to make a saccade to the fractal and fixate on it (500 ± 100 ms). Successful performance led to a large (0.3 ml) or small (0.1 ml) reward (33-66% diluted apple juice) based on the fractal identity (Good/Bad) concurrent with a correct tone. Fixation breaks or premature saccades to the fractal resulted in an error tone. Each trial was followed by an inter-trial interval (ITI) of 1-1.5s with a blank screen. The training block consisted of 80 trials (10 presentations/object) and objects presentation were shuffled pseudo-randomly.

### Familiarity Training

A random set of 8 fractals were chosen for perceptual familiarization for each monkey. Familiarization was done using passive viewing and free viewing over multiple days and sessions (>10 days). The details of both tasks are described below.

### Behavioral Training: Passive Viewing

Animal was required to keep a central fixation while familiar objects were displayed randomly with 400ms on and 400ms off schedule. Animal was rewarded for continued fixation after a random number of 2-4 objects were shown followed by 1-1.5s ITI. Objects were shown randomly in 8 radial directions with eccentricities from 5°-20°. A session of passive viewing with a set of eight familiar fractals had ∼20 presentations per object.

### Behavioral Training: Free Viewing

Each free viewing session consisted of 15 trials. In any given trial, four fractals would be randomly chosen from a set of four good and four bad objects in GB free viewing and from a set of four novel and four familiar objects in NF free viewing. For familiarization training all eight fractals were from the familiar set. Location and identity of fractals shown in a trial would be chosen at random in one of two possible configurations (for details see ^3^). The chosen fractals in a given trial were shown in any of the four corners of an imaginary diamond or square around center (15° eccentricities). Fractals were displayed for 3 sec during which the subjects could look at (or ignore) the displayed fractals. There was no behavioral outcome for free-viewing behavior. After 3 sec of viewing, the fractals disappeared. After a delay of 600 ± 100 ms, a white fixation dot appeared in one of nine random locations in the screen (center or eight radial directions). Monkeys were rewarded for fixating that fixation dot. The next trial was preceded by an ITI of ∼1.5 sec with a blank screen.

### Scanning: Passive Viewing

Each run consisted of 16 blocks lasting 30 sec each (eight base/eight probe). There were four different probe blocks: [good, bad] x [left, right] in GB scans and [novel, familiar] x [left, right] in NF scans (Fig. 1e). The order of four probe block types was pseudorandomized such that in every four sequential probe blocks all four types were shown and each run consisted of two such randomized cycles. Each block was divided into trials with variable duration (approximately four to five trials per block), which required central fixation on a 0.5° white dot within a 3°x3° window. During probe blocks, fractals were flashed (600 ms on, 200 ms off) in one hemifield (∼6° eccentricity) in horizontal and 45° oblique directions (pseudorandom location for each object) while the animal maintained central fixation. Three to five consecutive objects were shown during each trial. After the trial, central fixation was extinguished and the animal was rewarded (50–100% apple juice) for keeping fixation leading to an ITI of 2.2 sec. All objects in a given block were chosen from either good or bad categories in GB scans and novel and familiar categories in NF scans and were shown on either the right or left visual hemifields. No contingent reward for objects was delivered during passive viewing. Breaking fixation or failing to fixate centrally within 3 s of fixation dot appearance resulted in extinction of all visual stimuli followed by a 2.2 s ITI before start of the next trials. The data analyzed in this study consisted of 19 runs for monkey U and 18 runs for monkey D in GB scans and 20 runs for monkey U and 23 runs for monkey D in NF scans.

### fMRI Data Preprocessing

fMRI data is first converted form Bruker into AFNI file format. Slice time correction (3dTshift), motion correction (3dvolreg), and correction for static magnetic field inhomogeneities is performed on the data. After aligning the data with the anatomical FLASH images of each session, images were de-spiked, detrended (fourth-order Legendre polynomials), and transformed into percent change from the mean. The high-resolution anatomical images were transformed into standard D99 atlas space. For more details on preprocessing steps see ^21^.

### fMRI Analysis

The analysis was performed in AFNI and SUMA software packages (AFNI_18.2.09), Statistical Parametric Mapping (SPM 12), as well as custom-written MATLAB programs. The 3D+time series were regressed against the model time series to calculate the beta coefficients that represented the contribution of each factor in percent change from the mean (GLM analysis using 3dDeconvolve). The model time series consisted of eight regressors ([good, bad, novel, familiar]×[right, left hemifield]) that were one during the block satisfying their condition and zero otherwise and were then convolved with a MION hemodynamics. Seventeen nuisance regressors were used including motion and their first derivatives (12 parameters), reward delivery, blinks, and eye position (horizontal, vertical, and interaction). All nuisance time series except the ones related to motion were convolved with MION hemodynamics before regression (the negative change of MION agent was considered). A separate regression with one regressor (probe vs. base) and the same nuisance factors was also carried out to find visual beta coefficients for differential response to probe vs. base blocks. These beta coefficients reflected the degree of visual responsivity of voxels and were orthogonal to value-coding and spatial-coding voxels (i.e., switching the value or hemifield labels for a voxel does not change its overall activity). These coefficients were used to select the equal number of most visually active voxels across different anatomical regions (Fig. 2c-d). This selection method not only avoided the bias towards the number of selected voxels through each region, but also orthogonalized the selection to value-coding, novelty-coding, as well as spatial-coding. Blocks where subjects refused to fixate or were asleep were excluded from the analysis (<%10 monkey D and <11% in monkey U of all probe blocks and <6% in both monkeys base blocks). Also, the first three TRs in each run were excluded from the regression due to magnetization.

### Functional Connectivity Analysis

The residuals of the GLM analysis were used for resting state network extraction. In order to prevent common sources of correlation (confounders), the average white matter and ventricle time series were regressed out from this residual time series. The result was band passed between the desired frequencies (0.01–0.1 Hz) using the AFNI 3dBandpass function. The first base block period at the beginning of each run was selected and concatenated across all runs (excluding first three TRs for magnetization). In addition to NF and GB scans, first rest block from months-later GB scans were also used to increase statistical power (see ^21^ for details). ROIs for each anatomical region were selected based on union of significant NF and GB coding masks for each monkey (NF-GB mask) Regions with small number of significant voxels were omitted from graph analysis (<3-5 voxels). The weighted undirected graph was extracted from anatomical areas in the NF-GB mask using a constraint-based method for causal structure learning known as PC-stable algorithm. This algorithm uses a series of conditional independence tests (i.e. partial correlations) to extract the underlying Directed Acyclic Graph (DAG) associated with a given dataset. The significance level of the test which controls the sparsity of graph edges was a hyper parameter that was set manually for each monkey (1e-4 for Monkey D, 5e-6 for Monkey U). The algorithm guarantees extraction of the skeleton independent of the order of the conditional independence tests levels. Having extracted the graph skeleton, the smallest partial correlation associated with each pair of nodes was assigned as the weight of the corresponding edge.

Clustering of the derived weighted undirected graph was performed using spectral clustering algorithm on the largest connected components of the graph. The representation of the nodes in the space of Laplacian matrix eigenvectors lead to selection of clusters for each monkey. The clustering in the spectral space was performed using k-means algorithm with number of clusters set to four. These analysis steps are depicted in Fig 3a.

The functional connectivity of the subcortical substrates with each cluster centroid was evaluated through averaging all areas in each cluster together as the representative time-series of each cluster. Then correlation coefficient (Pearson’s) between all voxels of the four subcortical regions and the representative time series were obtained. The resulting correlations were thresholded based on the p-value and small-volume cluster-corrected to avoid false-positive clusters (P<0.001,).

### Dynamic Connectivity Analysis

The Dynamic Causal Modeling (DCM) algorithm was used to further analyze the novelty and value processing between the temporal and prefrontal nodes of the TP network. The method allowed us to evaluate the temporal dynamics during the task in the TP network. Hence the data used for this analysis consisted of the task fMRI, with seventeen nuisance factors regressed out (see fMRI Analysis for description of the nuisance factors). The DCM model considered here consisted of a neural state equation and a hemodynamic model. The neural state equation is formulated in equation 1.

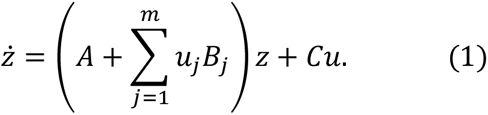

Where, state *Z* and inputs *u* are time-dependent and represent *k* nodes and *m* inputs of the model, respectively. *A* is a *k* × *k* input-independent connectivity matrix. *k* × *k* matrices (*B*_1_, *B*_2_, …, *B*_*m*_) were input-dependent modulations on the connectivity matrix by inputs (*u*_1_, *u*_2_, …, *u*_*m*_) respectively, and *k* × *m* input matrix *C* is the direct effect of inputs on the *k* nodes. In our case we had four nodes (k=4) and 8 different inputs (m=8, see below).

Four ROIs ([temporal, prefrontal] × [left, right hemisphere]) were selected based on the functional connectivity graph in Fig. 3. The mask was then used to extract the representative time-series of each node using the SPM volume of interest toolbox. A total of nine model configurations were investigated to arrive at the best of model. These models consisted of three different configurations for the input matrix multiplied by three different conditions for modulation matrix. In all models the fixed connection matrix was considered the same (supp Fig 7). In addition, we forced all models to satisfy the same matrix for both NF and GB tasks. The main question of interest, here, was addressing the differences and similarities of TP network connections regarding the task context and further analyzing the best model satisfying the well-known reciprocal connection between these regions. The competing architectures included 3 different input schemes with direct contralateral visual input to the temporal node (Temporal Input) to the prefrontal node (Prefrontal Input) and to both nodes (Double Input) (These were differences concerning input matrix in the model). On the other hand, as the question of interest was regarding the effect of the task type (NF or GB) on the connection of these two distinct cortical regions, we set a modulation of “Good” and “Novel” (left and right) stimuli on both forward and backward connections of temporal and prefrontal nodes in all models. In the first model (termed simple model) we observed a significant negative feedback from the prefrontal cortex to the temporal nodes in both monkeys. Given the need for network stability, the observed negativity may be due to stability requirements in the DCM model rather than real inhibitory role of prefrontal region. To further check this possibility, we allowed the modulation in matrix B to include self-connection terms (the diagonal entries of matrix). For each input configuration (“Temporal Input”, “Prefrontal Input”, and “Double Input”) we considered three variants with different modulations on self-connection of nodes. These connections were considered as free parameters to let the model stabilize itself without the need for negative feedback between the nodes. The first model named as “simple” had no modulation on the self-connections, in contrast the second and third models consisted of G/N (left and right) and G/B/N/F (left and right) self-modulation on all nodes respectively. Bayesian model selection included in the SPM package was used to choose among the nine competing models (3 input architecture x 3 self-connection modulation).

## Acknowledgements

We thank David Yu, Charles Zhu, and Frank Ye for assistance with fMRI scanning. This work was supported by the Intramural Research Program at the National Eye Institute. Functional and anatomical MRI scanning was carried out in the Neurophysiology Imaging Facility Core (National Institute of Mental Health, National Institute of Neurological Disorders and Stroke, and National Eye Institute).

## Authors Contributions

A.G., D.A.L., and O.H. designed research; A.G. and W.G. performed research; M.F., A.A. and A.G. contributed new reagents/analytic tools; M.F. and A.G. analyzed data; and A.G., M.F. D.A.L., and O.H. wrote the paper with input from other authors.

## Data and code availability

The data and analysis code used in this study are available upon reasonable request from the corresponding author.

---

**Supplementary Figure 1:**
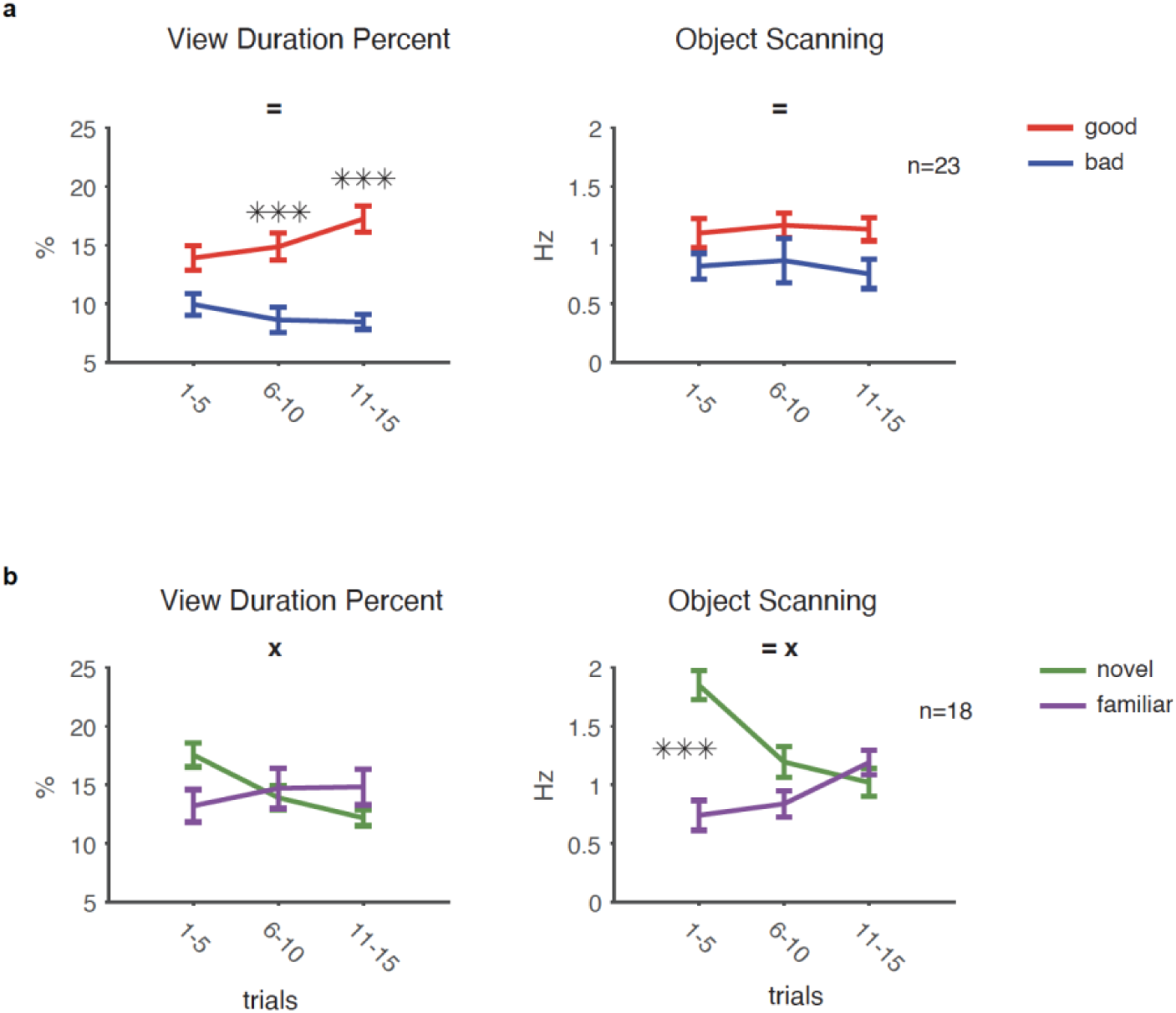
Sustained gaze bias for value dimension and transient gaze bias for novelty dimension. Objects were randomly selected and shown to the monkey for viewing in the absence of reward (GB and NF in separate blocks of free viewing) **a,** Gaze bias for viewing good vs bad object was measured during each session of free viewing and averaged separately for early (trials:1-5), mid (trials:6-10) and late (trials:11-15) epochs. (= significant main effect of value or novelty, \ significant main effect of trial, x significant interaction). Post-hoc tests (hsd) are marked with asterisks. Gaze bias measures shown are viewing duration as a percentage of total trial duration and the frequency of object scanning saccades while viewing the object. (F_1,132_>9.35, p<0.002 for main effect of value, F_2,132_<0.68, p>0.5 for main effect of trials, F_2,132_<2.86, p>0.06 interaction) **b,** same format as **a** but for novel vs familiar gaze bias. (F_2,102_>3.99, p<0.02 interaction and for object scanning F_1,102_=19.7, p<1e-3 main effect of novelty)

**Supplementary Figure 2:**
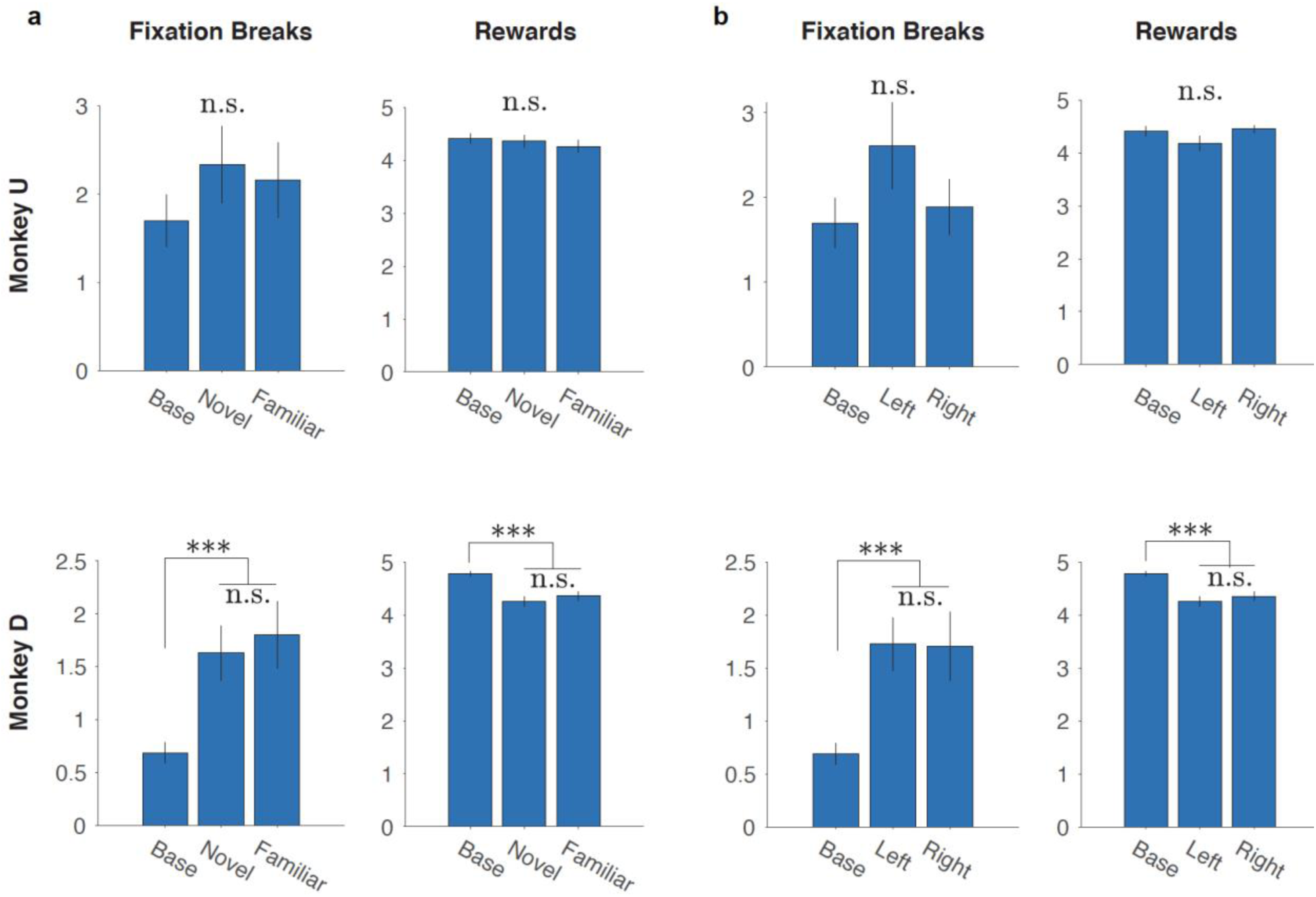
Equivalent performance and rewards across blocks during NF scans. **a,** Number of fixation breaks (left) and number of rewards received (right) in the base, novel and familiar presentation blocks for each monkey. (monkey U F_2,317_<0.89, p>0.41 and monkey D F_2,317_>10.4, p<1e-4, but difference between novel and familiar blocks not significant *hsd* post-hoc p>0.6) **b,** same format as in **a** but for the base, left, and right presentation blocks. (monkey U F_2,317_<1.69, p>0.18 and monkey D F_2,317_>10.3, p<1e-4, but difference between novel and familiar blocks not significant hsd post-hoc p>0.6)

**Supplementary Figure 3:**
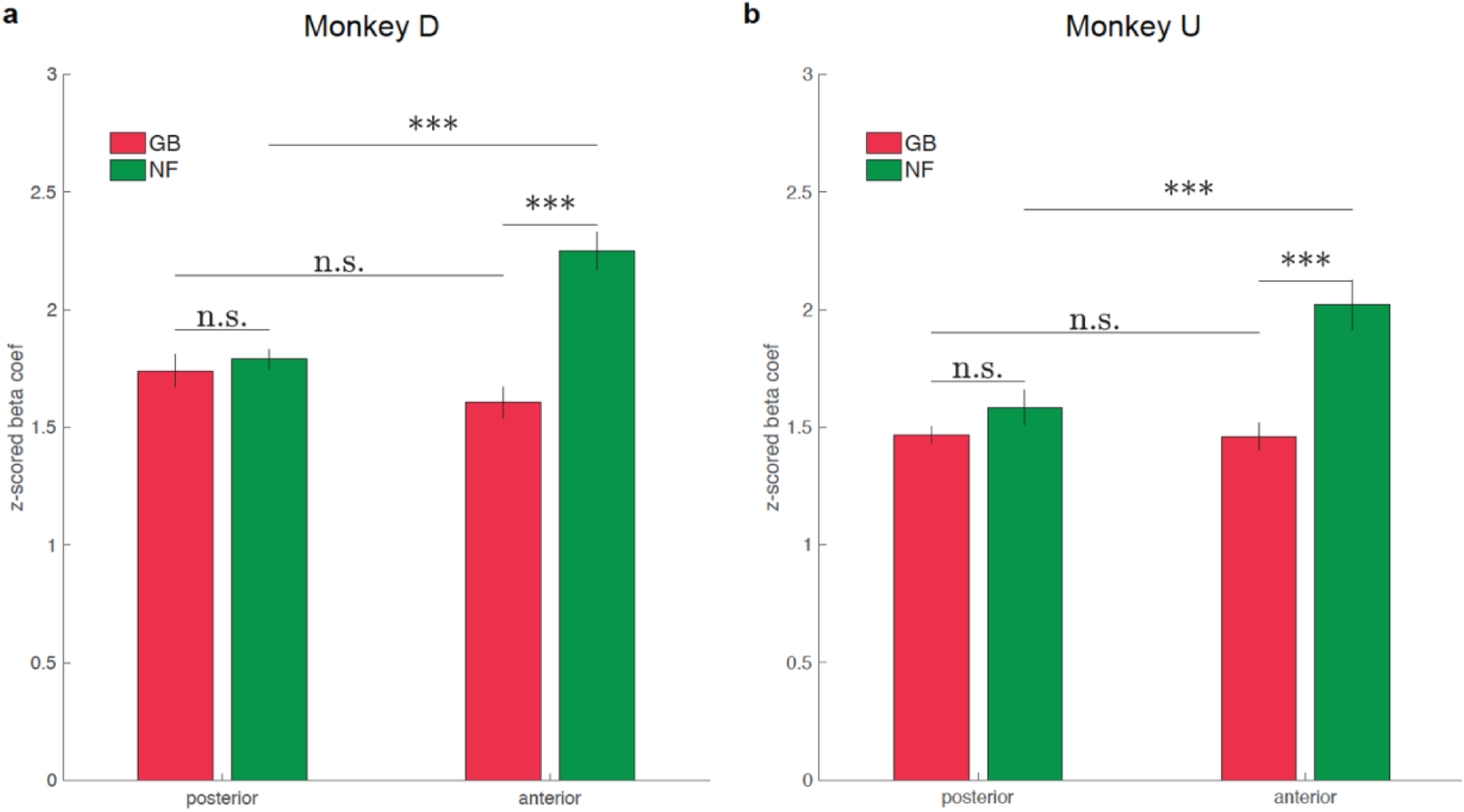
Difference of NF and GB coding in the anterior and posterior parts of vSTS. **a,** NF and GB beta coefficients z-scored using all gray matter beta coefficients to render the two dimensions comparable for monkey D. **b,** Same as **a** for monkey U. Anterior vSTS included TEm/a and posterior vSTS included TEO, FST, IPa, V4 and MT. Monkey U: F_3,522_ = 10.37, p = 1e-6, Monkey D: F_3,771_ = 10.50, p = 3e-8. Post-hoc test *hsd* ***P<0.001.

**Supplementary Figure 4:**
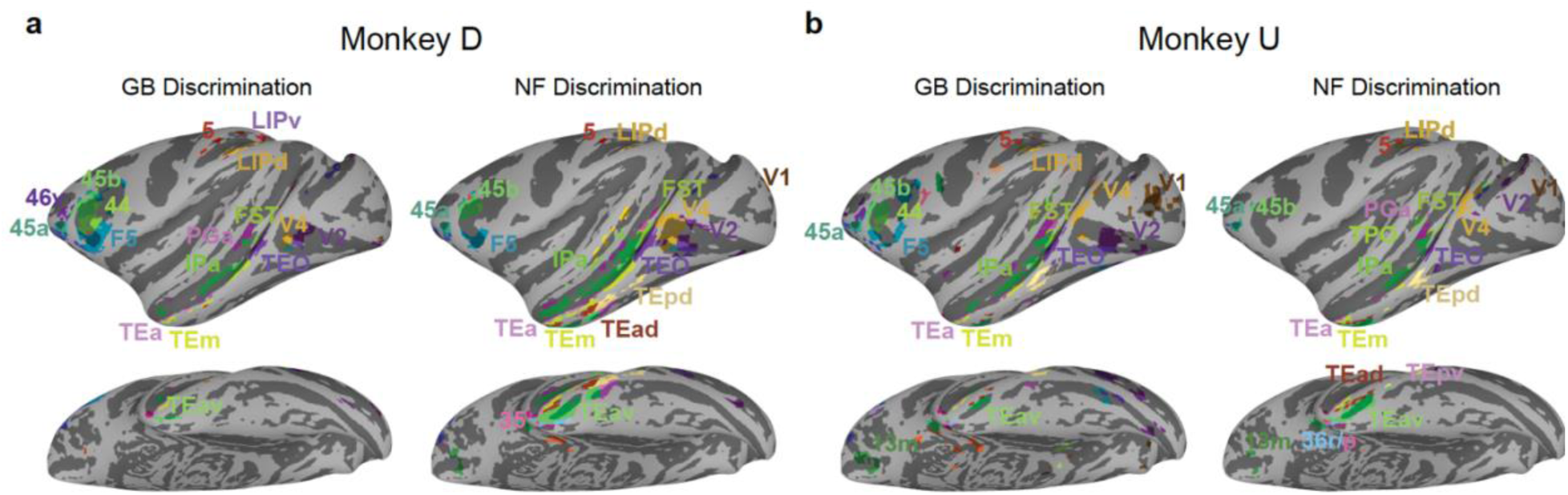
Annotation of cortical areas with significant GB and NF coding. **a,** Cortical areas corresponding to areas shown in Figure 2 in monkey D. **b,** Same format as **a** for monkey U.

**Supplementary Figure 5:**
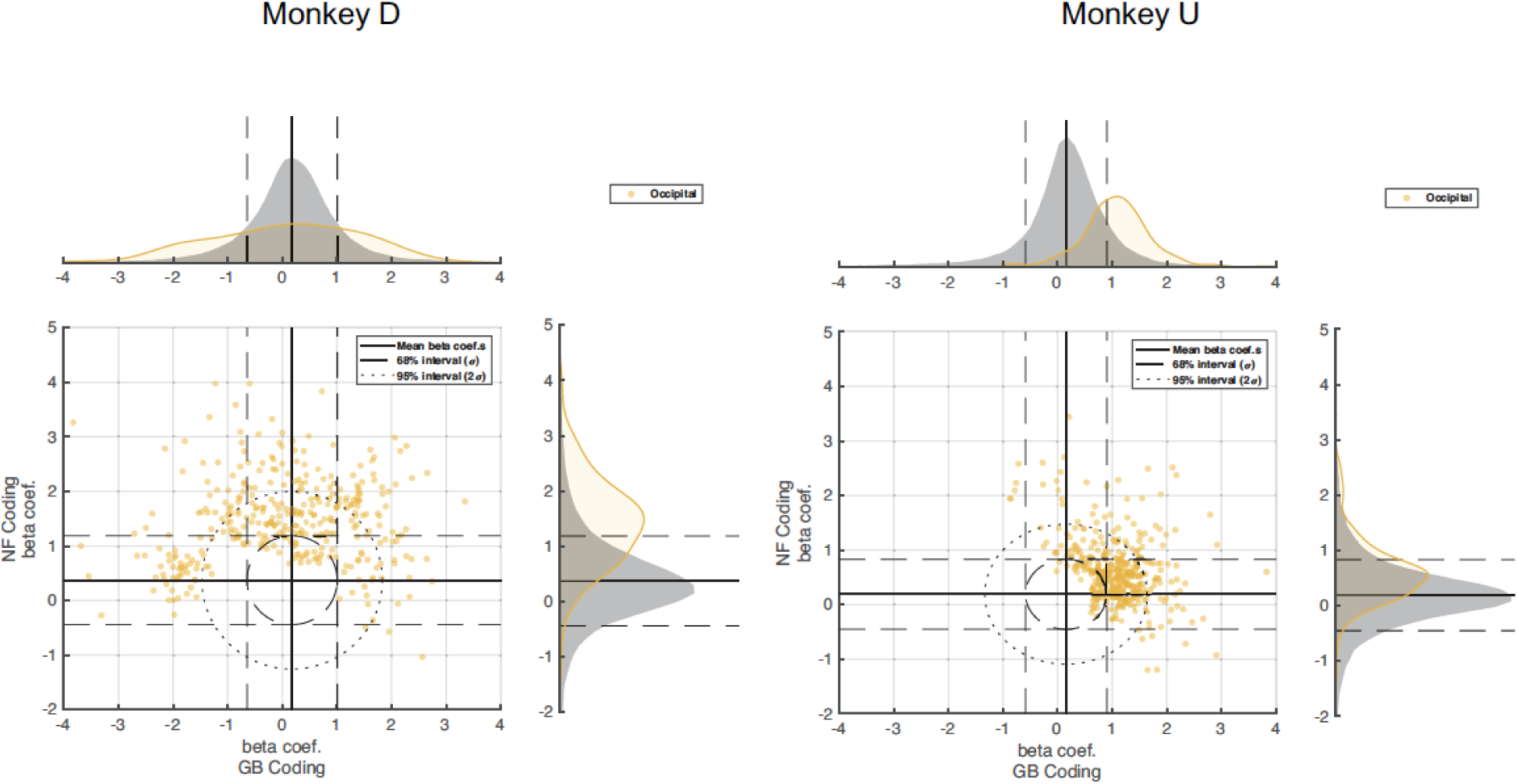
Novelty and value coding in the occipital network. Same as Figure 3b,**d** but for occipital network for both monkeys. All voxels are plotted without regards to the 68% and 95% thresholds.

**Supplementary Figure 6:**
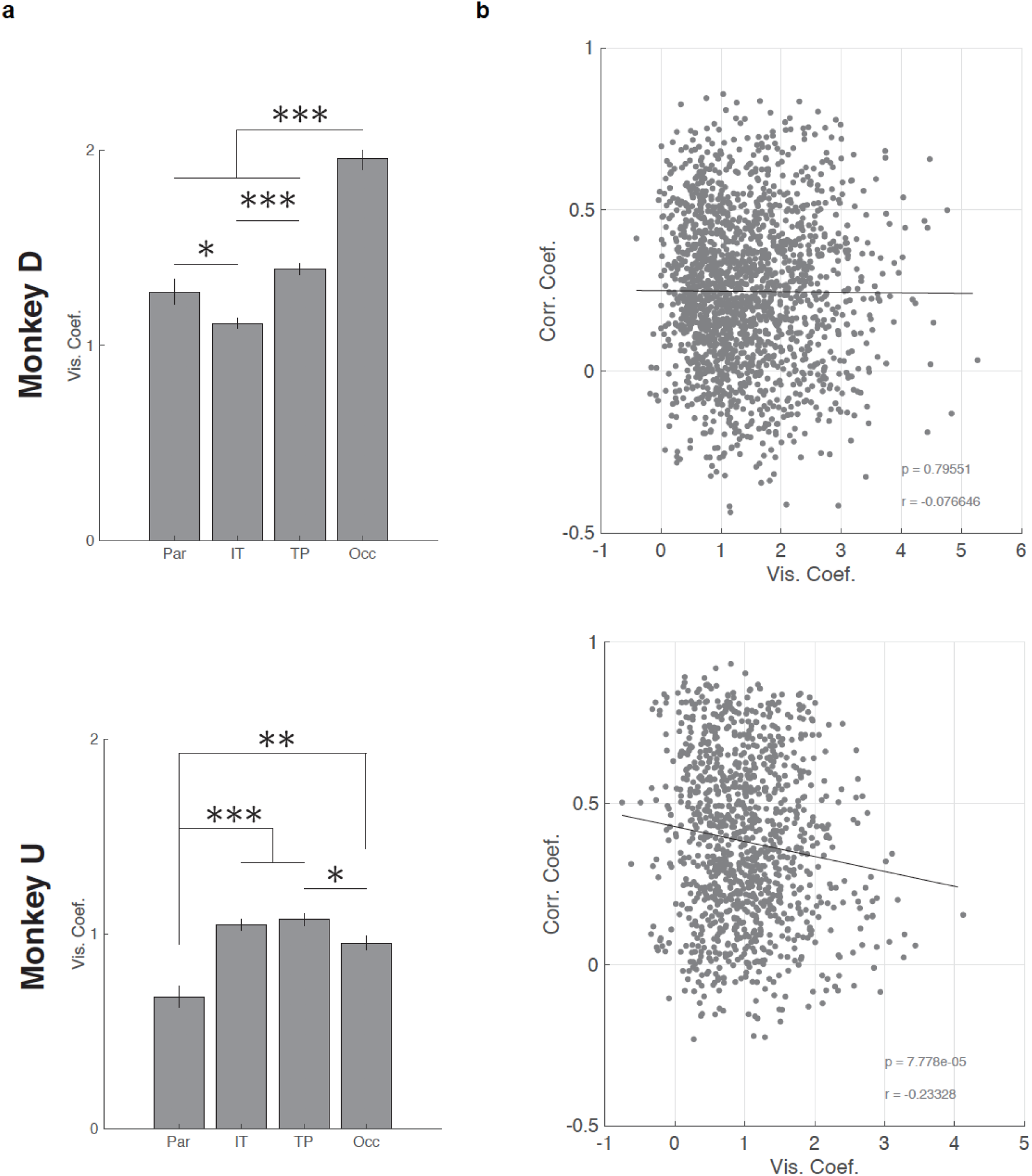
Visual activation vs NF/GB coding across cortical networks. **a,** The average visual coefficients across networks for each monkey. **b,** The relationship between the NF-GB time-series correlation and visual activation of all voxels in all networks (each point represents a voxel). r indicates Pearson’s correlation and the regression line is shown.

**Supplementary Figure 7:**
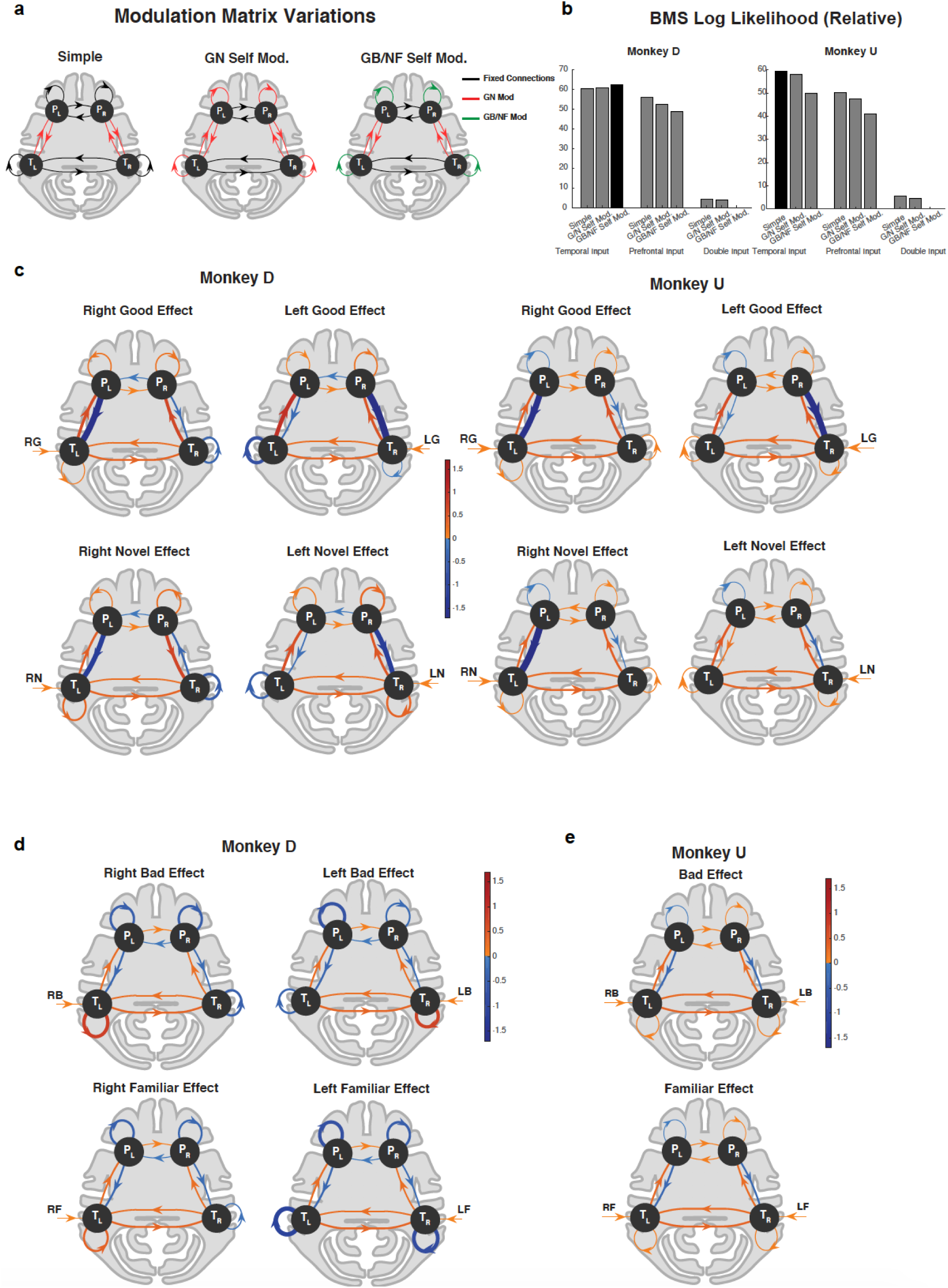
The DCM models considered for the TP network and the fitted weights for each object type. **a,** Three hypotheses regarding the modulation of self-connections. The black lines indicate no modulation. The red lines show edges modulated by the novel and good objects, and the green lines depict edges modulated by all object types: novel, familiar, good and bad. There were also three hypotheses about the input structure: only to temporal, only to prefrontal and to both structures (both) **b,** Bayesian model selection using relative log likelihood of nine proposed models for each monkey. In both monkeys the input only to temporal had higher likelihood. GBNF self-modulation was best for monkey D and no self-modulation (simple model) was best for monkey U. Best model with the highest posterior is colored black **c,** DCM model of the prefrontal-temporal interaction during good and novel objects separately for right and left blocks in both monkeys. (T_R,_ T_L_ :right and left temporal nodes, P_R,_ P_L_ :right and left prefrontal nodes). The line widths are scaled by the absolute value of the edges. Signed weights are color encoded. **d,** same as **c** but for bad and familiar object presentation for monkey D. **e,** same as **c** but for bad and familiar object presentation for monkey U. Since there was no modulation of weights by bad and familiar objects in this monkey both hemifield presentations shared the same weights.

**Supplementary Figure 8:**
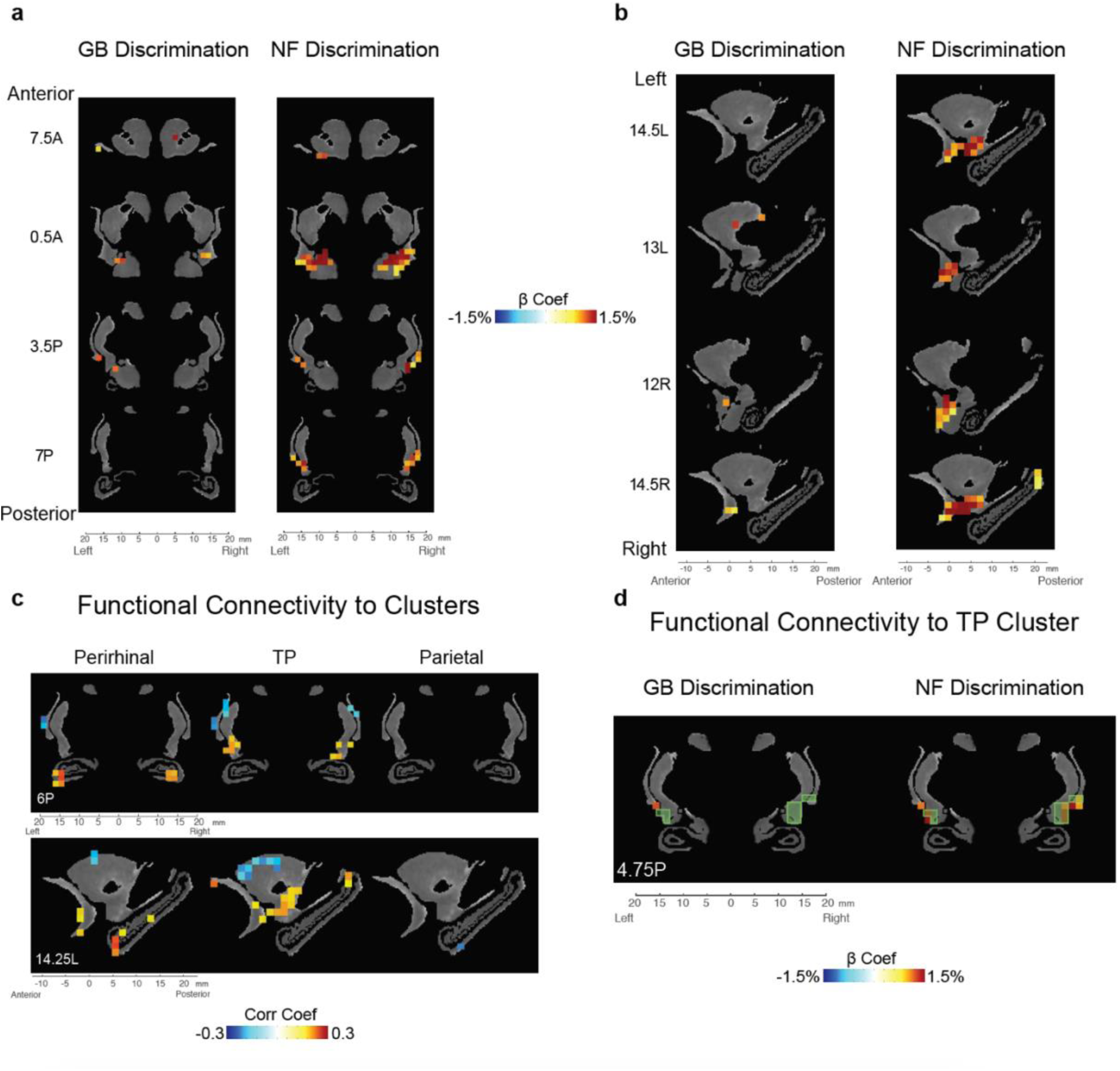
Subcortical novelty and value coding in striatum, amygdala, claustrum, and hippocampus. Same format as Figure 6 but for monkey D.

